# accuMUlate: A mutation caller designed for mutation accumulation experiments

**DOI:** 10.1101/182956

**Authors:** David J. Winter, Steven H. Wu, Abigail A. Howell, Ricardo B. R. Azevedo, Rebecca A. Zufall, Reed A. Cartwright

## Abstract

**Motivation:** Mutation accumulation (MA) is the most widely used method for directly studying the effects of mutation. Modern sequencing technologies have led to an increased interest in MA experiments. By sequencing whole genomes from MA lines, researchers can directly study the rate and molecular spectra of spontaneous mutations and use these results to understand how mutation contributes to biological processes. At present there is no software designed specifically for identifying mutations from MA lines. Studies that combine MA with whole genome sequencing use custom bioinformatic pipelines that implement heuristic rules to identify putative mutations.

**Results:** Here we describe accuMUlate, a program that is designed to detect mutations from MA experiments. accuMUlate implements a probabilistic model that reflects the design of a typical MA experiments while being flexible enough to accommodate properties unique to any particular experiment. For each putative mutation identified from this model accuMUlate calculates a set of summary statistics that can be used to filter sites that may be false positives. A companion tool, denominate, can be used to apply filtering rules based on these statistics to simulated mutations and thus identify the number of callable sites per sample.

**Availability:** Source code and releases available from https://github.com/dwinter/accuMUlate.

## Introduction

Mutations are important contributors to diverse processes such as adaptation, genome evolution, and the development of disease. Mutation accumulation (MA) is the classic method of directly studying the rates, molecular spectra, and fitness consequences of spontaneous mutations (Mukai, 1964). In a typical MA experiment, replicate inbred or clonal lines are isolated and repeatedly passed through severe bottlenecks. These bottlenecks reduce the effective population size of lines and thus reduce the efficiency of selection, allowing all but the most deleterious mutations to drift to fixation over the course of an experiment. The development of high-throughput sequencing technologies has led to a renewed interest in MA experiments. By sequencing whole genomes from MA lines, researchers can directly estimate the rate and molecular spectrum of mutations in a given species or strain. Studies combining MA experiments with whole-genome sequencing have provided key insights into the evolution of mutation rates (Long *et al.*, 2016; Lynch *et al.*, 2016; Sung *et al.*, 2012), genome evolution (Tenaillon *et al.*, 2016), the molecular basis of mutation (Ness *et al.*, 2015; Zhu *et al.*, 2014; Behringer and Hall, 2016), and the distribution of fitness effects among spontaneous mutations (Heilbron *et al.*, 2014; Dillon and Cooper, 2016).

The majority of the studies described above employ a custom bioinformatic pipeline to identify mutations from MA lines. In the widely used “consensus” approach (Ossowski *et al.*, 2010), a putative mutant is called if the majority of reads mapped to a given site differ from the most common base at that site across all samples. In an alternative approach, putative mutations can be identified by using variant calling software to call the most likely genotype for each MA line and the ancestral line at every site in the genome. In this approach, samples that are inferred to have a genotype they could not have inherited from the most-likely ancestral genotype are considered mutants (Zhu *et al.*, 2014). Because these approaches produce many false positive mutations, they are usually coupled with post-analysis filtering (based on sequencing coverage, the frequency of rare bases, or quality scores) to produce a final set of putative mutations. The particular filtering approaches used differ among studies. Putative mutations can then be validated by single-locus sequencing.

In this paper we describe accuMUlate, a mutation caller designed for MA experiments. Our approach can replace the custom pipelines and filtering processes currently used to analyze MA experiments with a unified approach to mutation calling. In addition to saving researchers time in developing custom pipelines, accuMUlate will increase the reproducibility of bioinformatic analyses of MA lines.

## Approach

accuMUlate uses the probabilistic approach to mutation detection described by Long *et al.* (2016). The probability that a given site in a genomic alignment contains at least one mutation is calculated from a model that directly reflects the design of MA experiments. In particular, we directly model the transition of alleles from an ancestral strain to descendant MA lines and account for the possibility of heterozygous sites in all lines. Our model also accommodates the noise associated with next generation sequencing data by using a Dirichlet-multinomial model to calculate genotype likelihoods (Wu et al., 2017).

For each putative mutation identified by accuMUlate, we calculate a suite of statistics that might be used to identify false-positive mutation calls (Li, 2014). In addition to statistics commonly used in existing approaches to mutation detection (including sequencing depth and mean mapping quality), this information includes the results of statistical tests for differences in quality control measures between sequencing reads containing apparently mutant bases and those that contain ancestral bases.

MA experiments are frequently undertaken in order to estimate the rate of mutation in a particular species, strain, or genotype. Accurate estimates of mutation rates require both a numerator (the number of sites at which a mutation was detected) and a denominator (the number of sites at which a mutation could have been detected if one were present). Our probabilistic approach to mutation calling provides a straight forward means to estimating this denominator. Mutations can be simulated for a given sample at a given site by altering bases in sequencing reads from that sample. Only those sites that generate a mutation probability greater than the threshold used for mutation detection and pass all additional filtering criteria used in a particular analysis should count towards the denominator for mutation rate calculations (Long *et al.*, 2016).

## Implementation

The accuMUlate package is written in C++ and contains two executable files. The main program, accuMUlate is a command line executable file written in C++. The program takes a single genomic alignment (in BAM format) with sequencing reads from the ancestral line (if they are available) and all MA lines as input. A number of additional arguments can be passed to accuMUlate to customize an analysis to a particular experiment. These arguments can be passed via the command line or through a simple text file. accuMUlate writes information for each site found to have a mutation probability higher than a user-set threshold. This information is provided in a tabular format, the first three columns of which correspond to the location of the putative mutation on the reference genome in standard BED format.

A second executable, denominate is included with the accuMUlate distribution. This program can be used to calculate the number of sites at which mutation could have been detected if one was present using the same parameters used in mutation calling and applying filtering criteria based on the statistics reported by accuMUlate. An additional repository, accuMUlate-tools (https://github.com/dwinter/accuMUlate-tools) provides a number of smaller utilities that may help users prepare files for accuMUlate and analyze the results.

The software is provided under an MIT license and is available from https://github.com/dwinter/accuMUlate/. The wiki at this repository (https://github.com/dwinter/accuMUlate/wiki) provides detailed instructions on how to compile the software, how to prepare files for analysis and how to interpret output files.

## Demonstration

We demonstrate the use of accuMUlate by reanalysing data generated from a previously published MA experiment. Shaw *et al.* (2000) allowled several lines of *Arabidopsis thaliana* to accumulate mutations. These lines have been the subject of two sequencing efforts. *Ossowski et al.* (2010) sequenced individuals from five lines to identify putative mutations, which they then validated by Sanger sequencing. Subsequently, Becker *et al.* (2011) generated longer sequencing reads from individuals representing 12 MA lines for an analysis of DNA methylation. We downloaded sequencing data from five of the lines Becker *et al.* (2011) sequenced, including two lines analysed by Ossowski *et al.* (2010). By comparing mutation calls generated by accuMUlate with the location of validated mutations (Wei *et al.*, 2014), we are able to demonstrate both the sensitivity of accuMUlate and the degree to which the various summary statistics reported for each site differ among validated mutations and other putative mutants.

A summary of this demonstration is described in a document provided as a Supplementary File. As desscribed in that supplement,accuMUlate, was able to recover all validated mutations along with a number of putative mutations that were not reported by *Ossowski et al.* (2010). The mapping-quality and insert-size statistics reported by accuMUlate differ substantially between validated and non-validated mutations. We were able to use these statistics to filter likely false-positives from all lines and, using denominate, to estimate the number of callable sites under these filtering criteria. Analysing this data further produces results similar to those reported by Ossowski *et al.* (2010). Our point-estimate of the mutation rate is slightly higher (8.3 × 10−^9^ base substitutions per site per generation compared 7 × 10−^9^ in the published work), but both stud show a mutational spectrum that is strongly biased toward G:C >A:T transitions. The source code for our demonstration is in the Supplementary Files for this paper.

These results show that accuMUlate accurately identifies mutations from MA experiments. The statistics reported for each putative mutation provide researchers with a straightforward way to detect potential false positives and denominate can generate a direct estimate of the number of callable sites in a given experiment. The reproducible data analysis in Supplementary File shows how the files produced by these programs can be used to produce the sorts of results usually reported from MA experiments. The accuMUlate distribution thus allows all of the key steps in the analysis of sequencing data from an MA experiment to be undertaken in a single framework.

